# Gut microbiota-derived GlcNAc-MurNAc is a TLR4 agonist that protects the host gut

**DOI:** 10.1101/2024.11.27.625627

**Authors:** Chenyu Li, Christopher Adamson, Allan Ng Wee Ren, Yaquan Liang, Zebin Hong, Jia Tong Loh, Siu Kin Ng, Jeric Kwan Mun Chung, Shiliu Feng, Evan Ng Wei Long, Nair Sajith Kumar, Christiane Ruedl, Sunny Hei Wong, Kong-Peng Lam, Yuan Qiao

**Author notes:** **Contact info** For correspondence.

## Abstract

Gut microbiota-derived peptidoglycan fragments (PGNs) are key signaling molecules that regulate multiple aspects of the host’s health. Yet the exact structures of natural PGNs in hosts have not been fully elucidated. Herein, we developed an LC-HRMS/MS analytical platform for global quantification and profiling of natural PGN subtypes in host gut and sera, unexpectedly revealing the abundance of PGN-derived saccharide moieties that do not resemble canonical ligands of mammalian NOD1/2 receptors. Focusing on the disaccharide GlcNAc-MurNAc (GM), a natural gut PGN that does not activate NOD1/2 yet still exhibits robust immunostimulatory effects in host immune cells, we unambiguously established GM as a TLR4 agonist, adding to the growing knowledge of NOD-independent mechanisms of PGN sensing in hosts. Importantly, the administration of GM effectively mitigates colonic inflammation in the DSS-induced colitis model in mice via TLR4-dependent mechanisms, highlighting the *in vivo* significance of natural gut microbiota-derived PGNs in maintaining host intestinal homeostasis.

## Introduction

The gut microbiota has a profound impact on host health. The resident gut bacteria bestow a rich source of microbial metabolites and molecules, known as microbe-associated molecular patterns (MAMPs), which are recognized by the host’s innate immune pattern recognition receptors (PRRs).^1,2^ Proper interactions between commensal bacterial MAMPs and host PRRs are critical for intestinal homeostasis, including the genesis and maturation of isolated lymphoid follicles (ILFs),^3^ reinforcement of gut barrier functions,^4,5^ and protection against colonic injury and pathogenic infections.^6,7^ Conversely, gut microbiota dysbiosis can inflict dysregulated immune responses in hosts, leading to chronic inflammatory diseases such as inflammatory bowel diseases (IBD), rheumatoid arthritis, and asthma.^8–11^ Understanding how the host recognizes and responds to gut microbial ligands is a key step toward developing novel immuno-therapeutics to target the gut microbiota-host interface.

Peptidoglycan, the major bacterial cell wall component, represents a well-known MAMP that stimulates the host’s innate immune system.^12,13^ A mesh-like layer that surrounds the bacterial cytoplasmic membrane, peptidoglycan is composed of repeating *N-*acetylglucosamine-*β*-1,4-*N*-acetylmuramic acid (GlcNAc-MurNAc) disaccharide with a stem peptide connected to the lactoyl group of each MurNAc unit. Adjacent peptidoglycan strands are cross-linked via the appended stem peptides.^14^ Despite a conserved scaffold, peptidoglycan varies significantly across bacterial species in terms of stem peptide composition, type and degree of cross-linking, and unique glycan modifications, giving rise to remarkably complex and heterogeneous peptidoglycan polymeric structures (*i.e.* peptidoglycome).^15^ During bacterial cell wall remodeling and turnover, soluble peptidoglycan fragments (PGNs) are generated and released into the milieu. These PGN fragments can activate mammalian NOD1/2 innate immune receptors, leading to the production of proinflammatory cytokines.^16^ Notably, NOD1 and NOD2 recognize distinct minimal PGN motifs: NOD1 detects the dipeptide D-γ-Glu-*m*DAP (iE-DAP),^17,18^ while NOD2 recognizes the muramyl dipeptide *N*-acetylmuramyl-L-Ala-D-isoGln (MDP or M-AQ).^19,20^ Biological studies of bacterial PGNs have predominantly focused on these canonical NOD1/2 ligands. Given the diversity of the gut microbiota peptidoglycome, these ligands likely cannot embody the repertoire of natural gut microbiota-derived PGNs in hosts, which may exhibit bioactivity via NOD-independent pathways. For instance, GlcNAc derived from peptidoglycan was shown to trigger NLRP3 inflammasome formation by inhibiting the metabolic enzyme hexokinase in primed immune cells without engaging NOD1/2.^21^ Moreover, we recently identified an anti-inflammatory 1,6-anhydro-PGN motif from probiotic *Bifidobacterium* that does not activate NOD1/2,^22^ suggesting alternate PGN sensing mechanisms in hosts. However, despite increasing recognition of gut microbiota-derived PGNs as key effector molecules in hosts, the fundamental questions regarding the structures of natural PGNs, their biological roles, and host responses beyond NOD1/2 signaling are yet to be determined.

Traditionally, detection of PGNs has largely relied on cell-based reporter assays that selectively respond to NOD1/2 ligands but not non-canonical motifs.^23^ We have developed a monoclonal antibody (mAb) 2E7 that specifically recognizes MDP, which enabled an indirect competitive enzyme-linked immunosorbent assay (icELISA) for PGN detection. This advancement has led to the discovery of gut microbiota-derived PGNs in host systemic circulation; however, the exact structures of natural PGNs remain unknown.^24^ Wheeler *et al.* recently employed radiolabeled ligands to elucidate the biodistribution of natural PGNs in hosts.^25^ Their research revealed that PGN-derived saccharide moieties preferentially accumulate in the host’s brain and adipose tissue, suggesting the physiological relevance of the non-canonical PGN motifs. Thus, it is imperative to uncover the natural forms of gut microbiota-derived PGNs to better understand their biological roles in hosts.

In this study, we developed an LC-HRMS/MS-based analytical platform to quantify and profile gut microbiota-derived PGNs in hosts. Contrary to the conventional notion, we did not identify canonical MDP (i.e. M-AQ) in the host gut; instead, we found that the PGN-derived saccharide moieties dominated the gut PGN pool. Unexpectedly, the PGN disaccharide motif, GlcNAc-MurNAc (GM), which does not activate NOD1/2, still robustly elicits immuno-stimulatory effects. Supported by cellular and biochemical evidence, we established that GM is a TLR4 agonist, highlighting the novel roles of PGN-TLR4 interactions in microbiota-host crosstalk. Importantly, we showed that GM administration effectively protects against DSS-induced colitis in mice via TLR4-dependent mechanisms. Our work underscores the biological significance of commensal gut bacteria-released PGN subtypes in safeguarding host intestinal homeostasis.

## Results

### Muramic acid (MurN) analysis affords global quantification of soluble PGNs in hosts

Acid hydrolysis of PGNs readily releases muramic acid (MurN), an invariant constituent of bacterial peptidoglycan that is absent in mammalian metabolites, rendering it an ideal unit for PGN quantificaition. Upon proper sample cleanup, we detected MurN (*m/z*: 252.1078) in both sera and feces of mice and humans (**Fig. 1A-C**), supporting the ubiquitous existence of gut microbiota-derived PGNs in hosts. Quantification of MurN was achieved based on the abundance of its two prominent daughter ions (*m/z*: 126.0549 and 144.0655) using the parallel reaction monitoring (PRM) mode of LC-HRMS/MS (**Fig. 1C**). Calibration curves were established with serial dilutions of *N*-acetyl-muramic acid (MurNAc) in water and spiked pooled sera or feces **(Extended Data Fig. 1).** Next, we quantified endogenous MurN in different biological samples with adjustments for the respective matrix effects. Ceca and feces from specific pathogen-free (SPF) mice contain a considerable amount of MurN, while feces from germ-free (GF) mice show virtually undetectable levels (**Fig. 1D**). Healthy human stools manifest higher MurN abundance but with considerable individual variability. Healthy human sera also display a wide range of MurN concentrations, averaging approximately 200 nM, which is comparable to the amount of MurN in two brands of fetal bovine serum (FBS). Notably, previous icELISA studies also revealed similar PGN concentrations in human sera,^24^ supporting the robustness and sensitivity of our quantification workflow. Together, our LC-HRMS/MS-based MurN analysis provides a quantitative perspective of gut microbiota-derived soluble PGNs in hosts.

**Figure 1.**
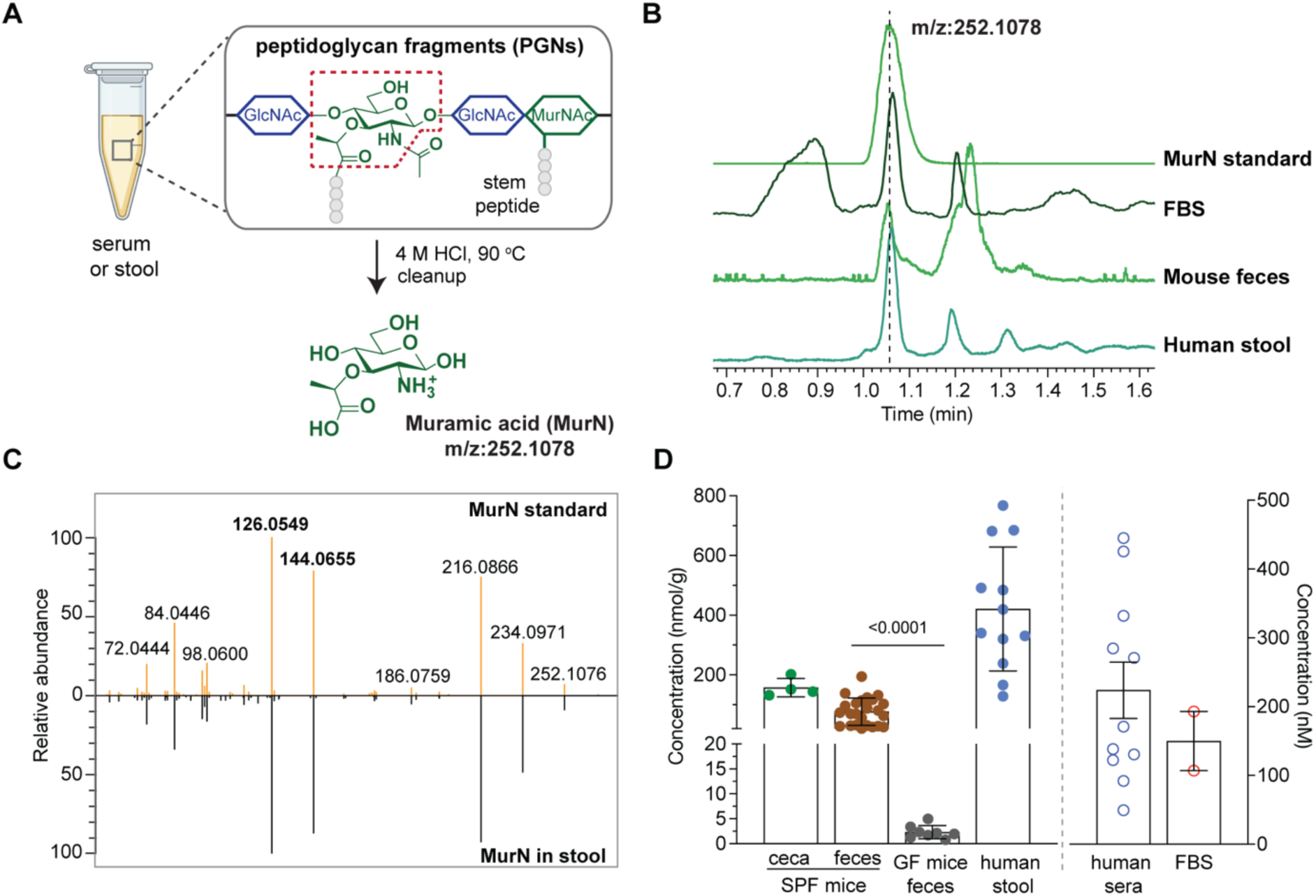
HPLC-MS/MS analysis of muramic acid (MurN) enables global quantification of gut microbiota-derived peptidoglycan fragments (PGNs) in biological samples. **(A)** Scheme of sample preparation for MurN analysis ([M+H]^+^: 252.1078). **(B-C)** Extracted ion chromatograms (EICs) and MS/MS spectra of MurN detected in host samples compared to the MurN standard. **(D)** MurN concentrations in biological samples, including SPF mice ceca (*n* = 4) and feces (*n* = 25), GF mice feces (*n* = 8), human stools (*n* = 12) and sera (*n* = 11), and commercial fetal bovine serum (FBS) from two different brands. Data are presented as mean values ± s.e.m. (*n* as indicated). Statistical significance was determined using one-way ANOVA.

### Saccharide moieties represent predominant PGN subtypes in the host gut

For structural profiling of gut microbiota-derived PGNs, we directly subjected the cleaned-up sera or stool samples to an untargeted analysis using the full-scan and data-dependent analysis (FSddA) mode of LC-MS/MS. Facilitated by our recently developed *in silico* PGN_MS2 spectral library,^22^ we extensively evaluated the experimental LC-MS/MS data for PGN identification, where hits were prioritized according to rigorous matching criteria including MS1 accuracy, isotopic pattern, and MS/MS fragmentation pattern (**Fig. 2A**). To validate potential PGN hits, we chemically synthesized a panel of PGNs as authentic standards (**Fig. 2B-D**). With this workflow, we found that the diversity and abundance of PGNs were significantly higher in the human stool than in sera, which is consistent with the rapid excretion and overall biodistribution of gut microbiota-derived PGNs in hosts.^26^ Interestingly, the major PGN subtypes in human stools and mouse feces are highly similar (**Extended Data Fig. 2**), suggesting that closely related gut bacteria may be the primary contributors of soluble PGNs, or that both mammalian hosts encode similar enzymes for processing PGNs.

**Figure 2.**
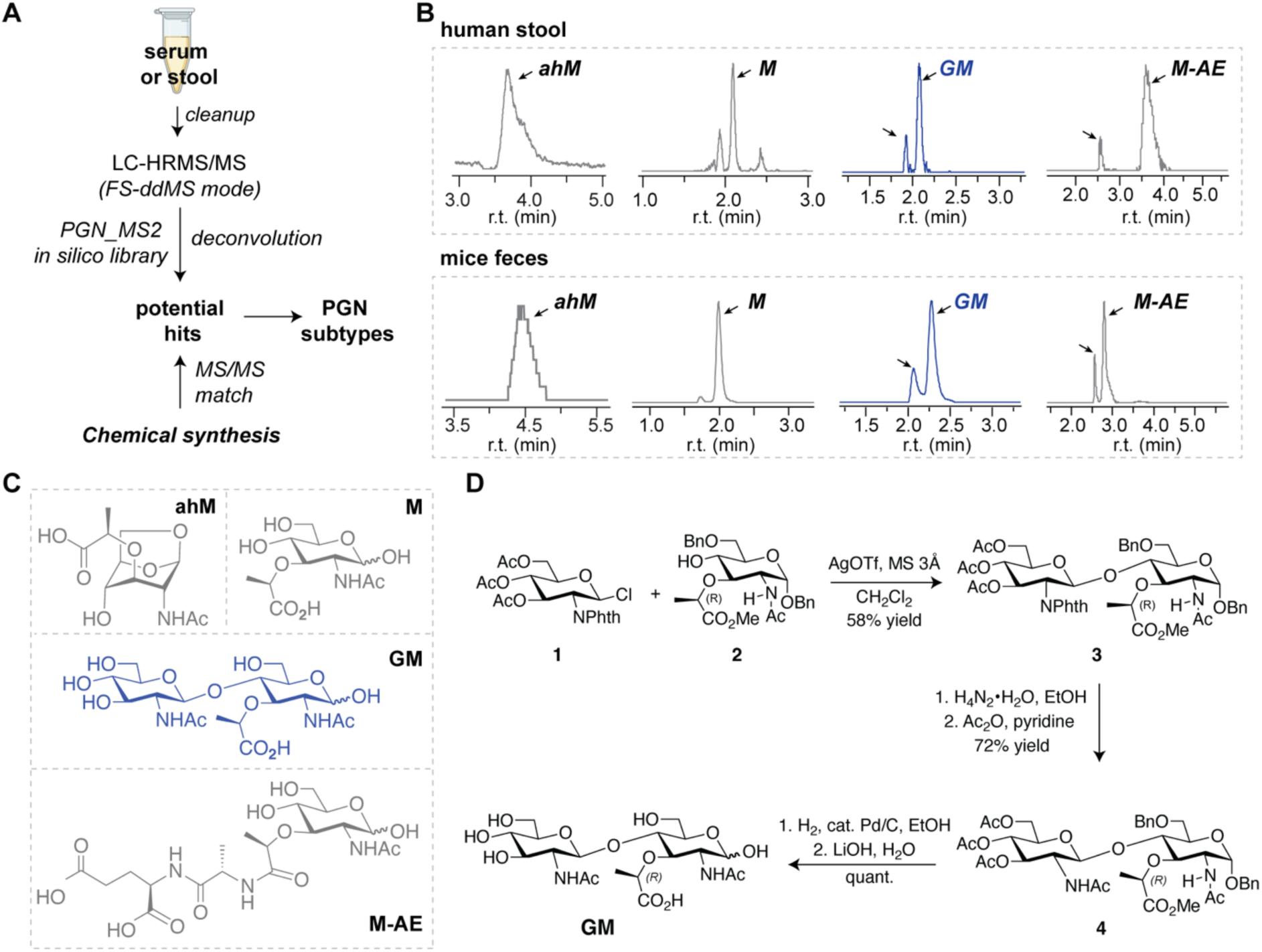
Profiling of gut microbiota-derived PGNs reveals abundant saccharide moieties in the host gut. **(A)** Scheme of HPLC-HRMS/MS workflow for PGN identification. **(B-C)** EICs of major PGN subtypes in the host feces: ahM, M, GM, and M-AE **(B)** and their chemical structures **(C)**. **(D)** Chemical synthetic route of disaccharide GM.

The soluble PGNs in the host gut can be classified into two groups: saccharide moieties that lack any stem peptide, and typical muropeptides containing both glycan backbone and stem peptide (**Fig. 2B-C, Extended Data Fig. 2**). The PGN-derived saccharide moieties, including MurNAc (M), 1,6-anhydro-MurNAc (ahM), GlcNAc-MurNAc (GM), and GlcNAc-ahMurNAc (G-ahM) are ubiquitously present in every sample of the cohort. In contrast, variable forms of muropeptides are detected across individuals, yet they mostly belong to the iE-DAP-containing PGNs. In addition, we revealed the abundance of muramyl-dipeptides GM-AE and

M-AE in mice ceca, which are close analogues of the widely used MDP (i.e. M-AQ) but differ in the amidation state of the stem peptide (**Fig. S1**). Furthermore, muropeptides with diacetylated GlcNAc or MurNAc moieties were detected, which represent peptidoglycan modifications that confer lysozyme resistance in many gut bacteria.^27^ LC-MS quantification of these major PGN subtypes showed that the saccharide moieties account for nearly 90% of all soluble PGNs in host feces. The diverse spectrum of natural PGNs in the host gut underscores the need to explore PGN bioactivities beyond the well-studied model ligands.

Profiling serum PGNs proved technically challenging due to low signal-to-noise ratios in LC-MS/MS analysis. Nonetheless, in two different cohorts of pooled sera from healthy human individuals and in commercial FBS, we identified PGN-derived monosaccharides M and ahM, but not any muropeptides (**Fig. S2**). The presence of M and ahM in serum coincides with their prominence as the two most abundant PGNs in the host gut (**Extended Data Fig. 2**), while the striking difference in the overall abundance and diversity of soluble PGNs between the gut and serum suggests that the systemic dissemination of gut microbiota-derived PGNs is a selective process.^25^ In a Caco-2 monolayer-coated transwell system, we showed that PGN saccharides M and GM translocated significantly faster than the larger muropeptide M-AEKAA (**Fig. S3A-B**). Furthermore, we reasoned that the absence of intact muropeptides in sera may be attributed to peptidoglycan-processing enzymes encoded by the host.^13,28–30^ For instance, mammalian PGRP2 is a serum amidase that specifically cleaves the *N*-acetyl-muramyl-L-Ala bond of soluble PGNs,^31^ whose activity may deplete muropeptides and contribute to the PGN-derived monosaccharides in host sera. Consistently, we showed that the incubation of 10% FBS with the muropeptide M-AEKAA reliably yields the expected degradation products: M and AEKAA (**Fig. S3C**). Taken together, our LC-MS/MS profiling workflow reveals that saccharide moieties are the predominant gut PGN subtypes in hosts.

### GM is immunologically bioactive via NOD-independent mechanisms

Intrigued by the abundance of gut microbiota-derived PGN saccharides in hosts at steady-state, we questioned if these natural subtypes exhibit any potential biological effects. To obtain sufficient PGNs of high purity for biological assays, we chemically synthesized a panel of PGNs of interest, including M-AE, GM, and M (**Fig. 2C-D**). As mammalian NOD1/2 represents canonical PGN sensors, we first assessed these PGNs in HEK-Blue^TM^ NOD reporter cells (**Fig. 3A**). As expected, the natural muropeptide M-AE potently elicited NOD2 activation similar to the model ligand MDP (i.e. M-AQ); yet the saccharide moieties, such as GM and M, are unable to activate NOD1/2, which is unsurprising given their lack of the required structural motifs for NOD1/2 recognition.^20^ At this juncture, we sought to explore the potential effects of PGN saccharides in immune cells that express diverse PRRs, which are more physiologically relevant than the NOD reporter cells. Unexpectedly, the disaccharide GM exhibits dose-dependent immuno-stimulatory activity in a range of immune cells, including murine macrophage RAW264.7, primary murine bone-marrow-derived macrophages (BMDM), and human monocytes (THP-1), as evidenced by the robust expression and production of proinflammatory cytokines such as TNFα and IL1β (**Fig. 3B-D**). In comparison, the monosaccharide M appears less potent (**Fig. 3C-D**), suggesting that the disaccharide structure may be crucial for immunoactivity. Furthermore, GM upregulates CD80/86 surface marker expressions in the type-II conventional DC (cDC2) population of murine BMDC (**Fig. 3E and S4**). Its ability to promote dendritic cell maturation *ex vivo* suggests a potential role in regulating adaptive immune responses in hosts. Importantly, we confirmed that the synthetic GM is endotoxin-free, as it tested negative in the LAL assay and its immunological effect unaffected by polymyxin B neutralization (**Fig. 3F-G, Extended Data Fig. 3**). In addition, no glycosidase-cleaved monosaccharides (G and M) were detected in GM-treated host cells, indicating that the immunoactivity likely originates from the intact disaccharide GM rather than its degradation products (**Extended Data Fig. 4**). With these observations, we concluded that the disaccharide GM is a bioactive PGN motif that acts via NOD1/2-independent pathways.

**Figure 3.**
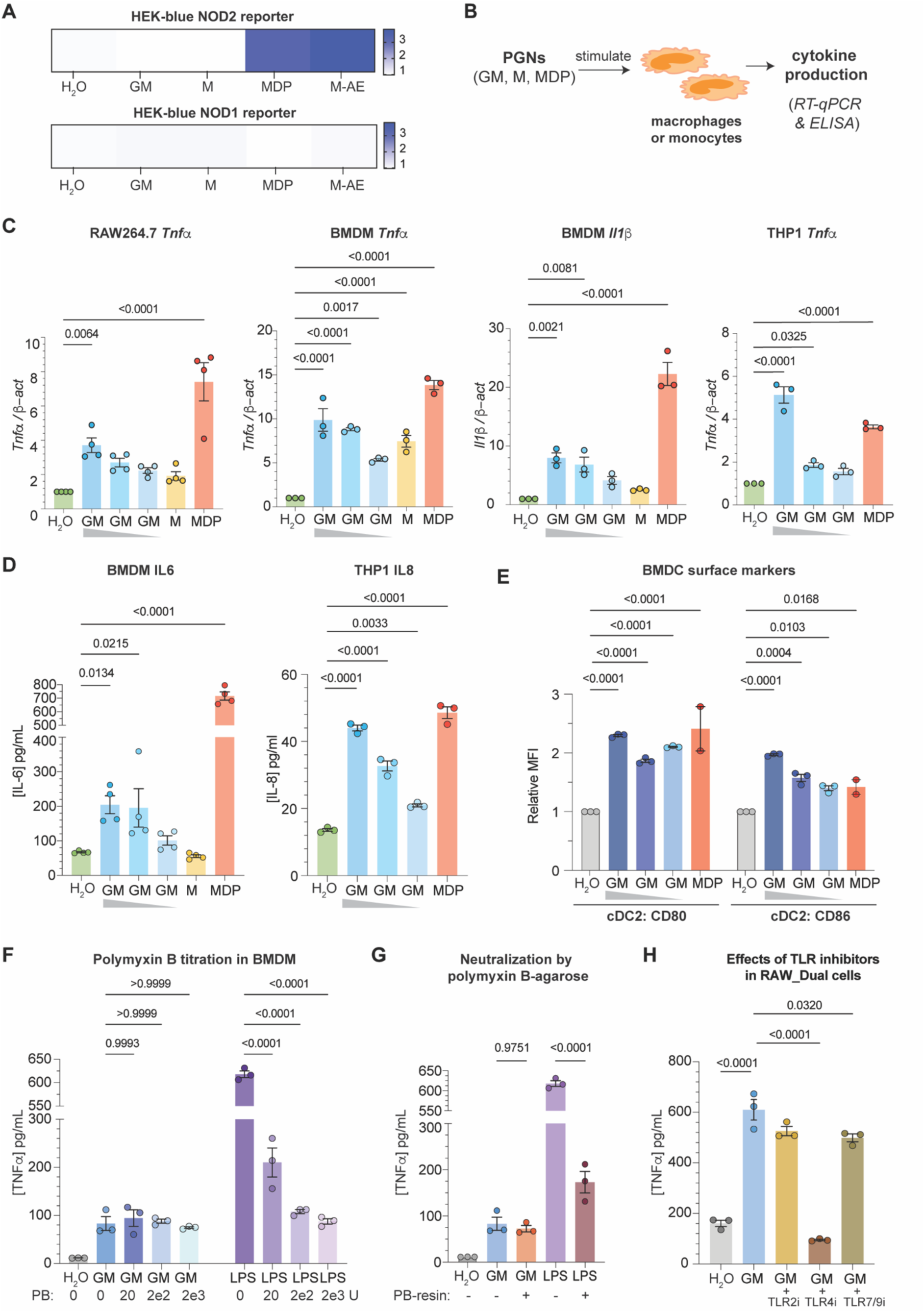
The disaccharide GM exhibits immuno-stimulatory effects *in vitro* independent of NOD1/2 receptors. **(A)** Readouts from HEK-Blue^TM^ NOD1/2 reporter cells stimulated with MDP (200 nM), M-AE (200 nM), GM (20 *μ*M), or M (20 *μ*M) for 18 h. Data are presented as mean values of biological replicates (*n* = 3). **(B-G)** PGNs induce cytokine production in macrophages and monocytes and activate dendritic cells. The respective immune cells were stimulated with the indicated PGNs for 24 h followed by cytokine analysis by RT-qPCR **(C)** or ELISA **(D)**. RAW264.7 cells were treated with GM (4 mM, 2 mM, 1 mM), M (4 mM), or MDP (0.4 mM); murine bone marrow-derived macrophages (BMDMs) and THP-1 cells were treated with GM (8 mM, 4 mM, 2 mM), M (8 mM) or MDP (0.4 mM). Data are presented as mean values ± s.e.m. of biological replicates (*n* = 3-4). Statistical significance was calculated using one-way ANOVA. (**E**) Mean fluorescent intensity (MFI) of surface CD80/86 in cDC2s population of murine bone marrow-derived dendritic cells (BMDMs) that were treated with MDP (20 *μ*M) or GM (50, 100, 200 *μ*M) for 18 h. Data are presented as mean values ± s.e.m. of biological replicates (*n* = 2-3). Statistical significance was calculated using two-way ANOVA. (**F-G**) Assays using Polymyxin B confirmed that GM is not contaminated with a trace amount of endotoxin. ELISA analysis of the TNFα levels in BMDMs stimulated with GM (4 mM) or LPS (100 ng/mL) titrated with Polymyxin B (0, 20 U/mL, 200 U/mL, 2000 U/mL) **(F)** or with Polymyxin B-neutralized GM or LPS **(G)** for 24 h. Data are presented as mean values ± s.e.m. of biological replicates (*n* = 3). Statistical significance was calculated using one-way ANOVA. (**H**) The TLR4 inhibitor specifically suppresses the GM-induced TNF-*α* production. The RAW_Dual^TM^ cells were pre-incubated with the respective TLR inhibitors (10 μM) for 1 h followed by GM stimulation (2 mM) for 24 h. TLR2i, TLR4i, and TLR7/9i refer to CU_CPT22, TAK-242, and AT791, respectively. Data are presented as mean values ± s.e.m. of biological replicates (*n* = 3). Statistical significance was determined using one-way ANOVA.

### TLR4 is essential for GM-triggered immune responses

Apart from the NOD sensors, toll-like receptors (TLRs) represent another important class of mammalian PRRs that detect various MAMPs, many of which contain glycan motifs.^32^ To determine if GM is recognized by TLRs in immune cells, we pretreated RAW264.7 cells with a panel of small-molecule TLR inhibitors before stimulating with GM, and then measured cytokine levels in the culture supernatant using ELISA. Remarkably, the addition of TAK-242, a potent and selective TLR4 inhibitor,^33^ drastically suppressed GM-triggered TNFα production in RAW264.7 cells, while inhibition of TLR2 or TLR7/9 did not interfere with GM’s bioactivity (**Fig. 3H and S5**). Our results implicate TLR4, a well-recognized innate immune receptor of bacterial LPS,^34^ in sensing the gut microbiota-derived disaccharide GM. Importantly, we demonstrated that TNFα production in GM-triggered BMDMs is unaffected by polymyxin B titration, which effectively sequesters LPS/endotoxin (**Fig. 3F-G**). This confirms that the observed TLR4-dependent bioactivity of GM is not due to trace endotoxin contaminants.

To conclusively establish the role of TLR4 in the immunostimulatory effects of GM, we next utilized RAW-Dual^TM^ wildtype (WT) and *Tlr4^-/-^* reporter cells for assays. Notably, TLR4 activation is known to trigger two distinct signaling transduction pathways, NF-*κ*B- and IRF-dependent pathways,^35–37^ both of which can be expeditiously evaluated in parallel via colorimetric and luciferase readouts in the dual reporter cells (**Fig. 4A**). As expected, GM induces dose-dependent activation of both NF-κB and IRF signaling in WT cells but exhibits minimal activity in *Tlr4^-/-^*cells, which supports the essential role of TLR4 in GM-induced immune responses. Interestingly, the regio-isomer MurNAc-GlcNAc (MG) shows similar TLR4-dependent immune activation (**Fig. 4B**).

**Figure 4.**
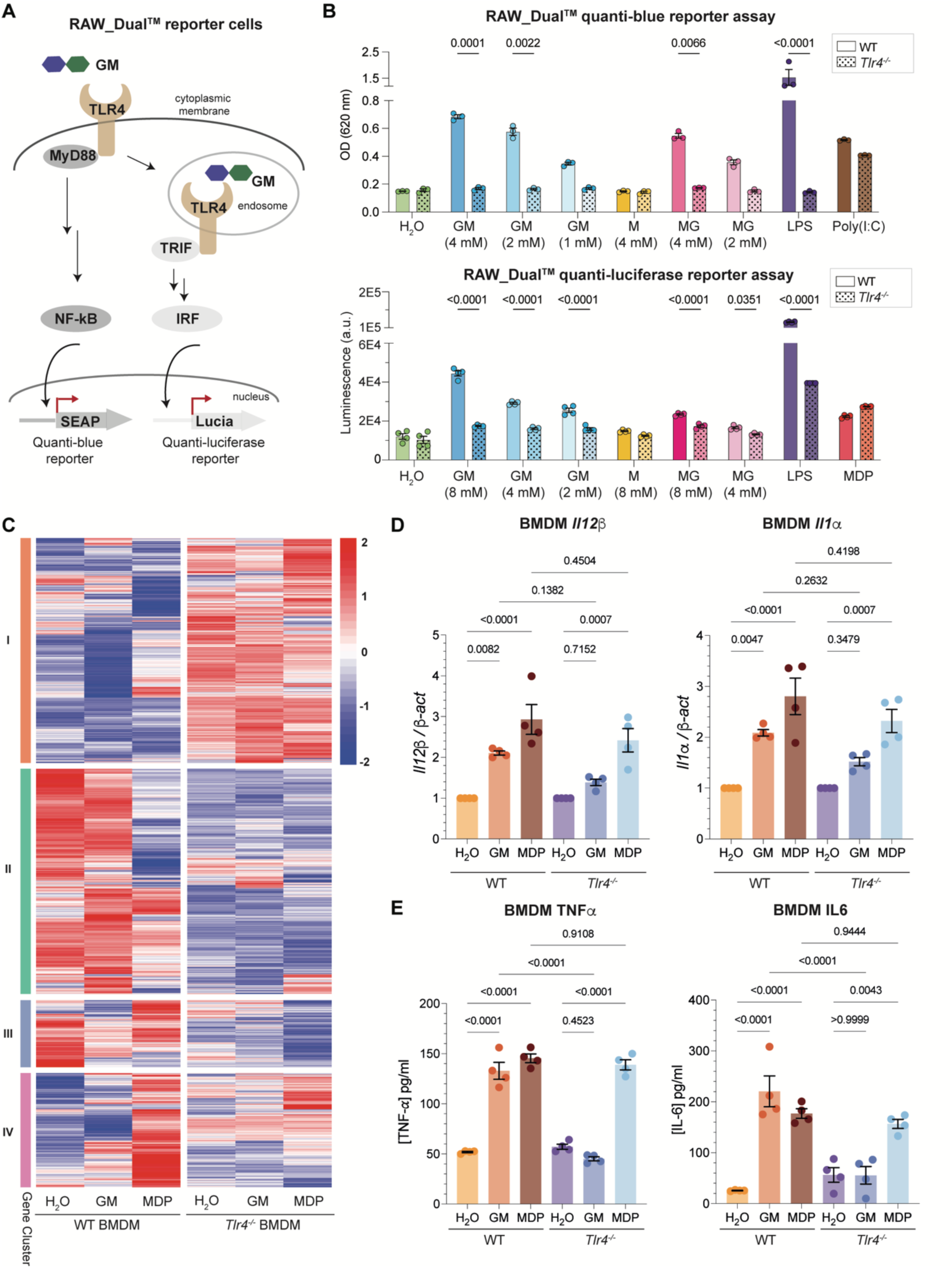
The immuno-stimulatory activity of GM is TLR4-dependent. **(A-B)** WT and *Tlr4*^-/-^ RAW_Dual^TM^ reporter cells were used to evaluate GM-induced NF-*κ*B and IRF signaling. Cells were treated with respective PGNs at indicated concentrations for 24 h for quanti-blue analysis (top) or quanti-luciferase analysis (bottom). LPS (100 ng/mL), MDP (0.4 mM), or Poly(I:C) (0.1 mg/mL) were used as controls. Data are presented as mean values ± s.e.m. of biological replicates (*n* = 3). Statistical significance was determined using two-way ANOVA. **(C-E)** The RNAseq analysis and validation confirm the GM-induced immune responses in BMDM require TLR4. **(C)** Heatmap plot of RNAseq results showing the mean values of differentially expressed genes (p < 0.01, and logFC > 1) in GM- or MDP-treated WT and *Tlr4*^-/-^ BMDMs (*n* = 3 biological replicates for each group). WT and *Tlr4*^-/-^ BMDMs were treated with GM (8 mM) or MDP (0.4 mM) for 24 h. (**D-E**) Validation of the RNAseq results by RT-qPCR and ELISA. Data are presented as mean values ± s.e.m. of biological replicates (*n* = 4). Statistical significance was determined using one-way ANOVA.

To gain deeper insights into the GM-induced transcriptional effects that are TLR4-dependent, we performed whole-genome RNA sequencing (RNAseq) in WT and *Tlr4^-/-^* BMDMs that were stimulated with GM for 24 h. Gene expression levels were analyzed with DESeq2 (**Figure 4C**).^38^ KEGG pathway analysis of the GM-treated WT BMDM (versus H_2_O-treated cells) identified several upregulated pathways associated with immune response and autoimmune diseases, such as Toll-like receptor signaling, NF-kB signaling pathway, and inflammatory bowel diseases, which are not enriched in the GM-treated *Tlr4^-/-^* BMDMs (**Fig. S6A-B**). Specifically, a large number of cytokine and chemokine genes, including *Il12α/β*, *Il6*, *Il1α/β*, *Csf3*, *Ccl5*, and *Cxcl1*, are significantly upregulated in GM-treated WT BMDMs compared to the H_2_O-treated controls. In contrast, GM stimulation does not induce the expression of these genes in *Tlr4^-/-^* BMDMs (**Extended Data Fig. 5**). The transcriptional changes were further validated by RT-qPCR (**Fig. 4D, Extended Data Fig. 6A)**. Collectively, our results confirm that TLR4 is indispensable in mediating the immunological effects of gut microbiota-derived disaccharide GM.

Additionally, we also analyzed RNA-seq data from WT and *Tlr4^-/-^* BMDMs treated with MDP, the model NOD2 ligand (**Fig. 4C**). As expected, MDP-induced pathways---including TNF signaling, NOD-like receptor signaling, and NF-κB signaling---are similarly enriched in WT and *Tlr4^-/-^* BMDMs (**Fig. S6C-D**). Upon closer analysis of cytokines and chemokines, we observed that while many genes, such as *Il12* and *Il6*, are highly stimulated by MDP in both types of cells, a subset of MDP-induced genes, including *Ccl5*, *Cxcl10*, and *Tnfα*, appear to be TLR4-dependent (**Fig. 4D, Extended Data Fig. 5-6)**. This intriguing discovery suggests that TLR4 may also play a role in modulating MDP-triggered NOD2 signaling, which was previously unrecognized.

Lastly, we also evaluated cytokine production in treated *Tlr4^-/-^*BMDMs by ELISA. The disaccharide GM, like LPS, fails to elicit any significant level of TNFα and IL6 in *Tlr4^-/-^* BMDMs; in contrast, other ligands such as CpG and MDP still robustly trigger proinflammatory cytokine production in *Tlr4^-/-^* BMDM, similar to their effects in WT BMDM (**Fig. 4E, Extended Data Fig. 6B-D**). Of note, the lack of immune-stimulatory response in GM-treated *Tlr4^-/-^* BMDMs was paralleled with the complete absence of p65 and p38 phosphorylation, which was in contrast to the mild but robust increase in phosphorylation in GM-treated WT BMDMs (**Extended Data Fig. 7**).

### The disaccharide GM directly binds to TLR4

To conclusively establish TLR4 as the putative receptor for the disaccharide GM, we sought biochemical evidence of GM binding to TLR4. First, in an *in vitro* pulldown assay, we demonstrated that the immobilized GM-1 (GM-1-resin) effectively enriched recombinant mTLR4 protein in a dose-dependent manner, whereas the control resin immobilized with GlcNAc (G-1-resin) did not exhibit specific interaction with mTLR4. Pre-incubation of mTLR4 with excess GM as competitors led to a dose-dependent decrease in the amount of mTLR4 bound to GM-1-resin (**Fig. 5A-B**). To solicit additional evidence for direct GM-mTLR4 interaction, we next performed the surface plasmon resonance (SPR) assay (**Extended Data Fig. 8**), in which LPS binds to mTLR4 in a dose-dependent manner, while MDP does not interact with it, serving as appropriate positive and negative control ligands that validate the robustness of the SPR setup. As anticipated, the disaccharide GM binds to mTLR4 in the SPR assay, with an estimated Kd of ∼383 µM. The low binding affinity of GM to mTLR4 *in vitro* is consistent with its mild immunostimulatory effects observed in the cellular assays. Our data confirms GM is a TLR4 ligand.

**Figure 5.**
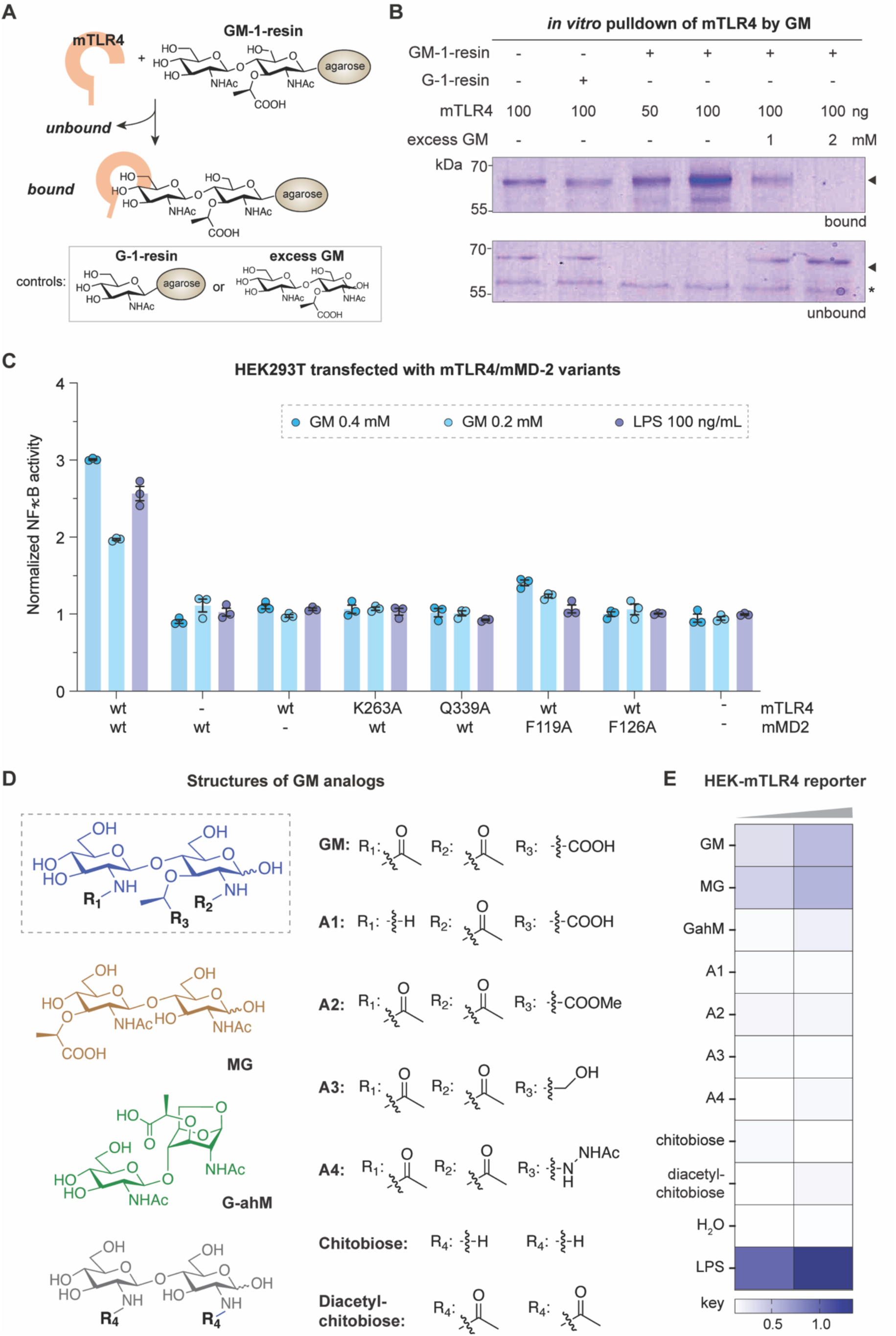
TLR4/MD-2 specifically recognizes GM. **(A-B)** *In vitro* enrichment of recombinant mTLR4 protein by immobilized GM. **(A)** Scheme of the pulldown assay, with GM- or G-immobilized agarose beads. **(B)** Coomassie blue-stained SDS-PAGE analysis of mTLR4 proteins bound or unbound to the resin. Excess GM was added to compete off mTLR4 bound to GM-immobilized beads. **(C)** Analysis of specific residues in TLR4/MD-2 required for GM-induced NF-*κ*B activity. The various mTLR4/mMD2 variants were transiently expressed in HEK-Blue^TM^ reporter cells and treated with GM (0.4 mM, 0.2 mM) or LPS (100 ng/mL) for 24 h. The activation of NF-*κ*B was measured based on the SEAP reporter signals. For each transfected group, the readouts were normalized to the H_2_O-treated samples. Data are presented as mean values ± s.e.m. of biological replicates (*n* = 3). (**D-E**) Structural-activity relationship (SAR) analysis of disaccharide analogues using HEK-Blue^TM^ mTLR4 reporter cells. Cells were treated with respective disaccharides at 0.5 mM or 1 mM. LPS (20 ng/mL and 100 ng/mL) was used as controls. Data are presented as mean values of biological replicates (*n* = 3).

### MD-2 is essential for TLR4 recognition of disaccharide GM

Of note, MD-2 is an essential co-receptor of TLR4 for cell surface sensing of LPS,^39^ where specific residues in TLR4/MD-2 are known to stabilize the hydrophobic lipid chains and the polar glycan core of lipid A.^40^ To investigate if MD-2 is also required for TLR4 recognition of GM, we evaluated the NF-*κ*B activity in GM-stimulated HEK293T cells that transiently express either mTLR4, mMD-2, or both, where the expression levels of the transfected genes were validated by Western blot (**Fig. 5C, Extended Data Fig. S9A-B**). Similar to LPS, GM exhibits dose-dependent activation of NF-*κ*B signaling only in cells that co-express mTLR4 and mMD-2, indicating MD-2 is essential for GM recognition by TLR4. Considering GM potentially resembles the disaccharide core of Lipid A, we predict that the TLR4/MD-2 residues involved in polar interactions are likely crucial for GM recognition. As expected, mutations of mTLR4_K263 and Q339, both of which form hydrogen bonds with the glycan core of Lipid A,^40,41^ effectively abolished GM-triggered NF-*κ*B activity; whereas mutating mMD-2_F119, a residue that stabilizes the hydrophobic acyl chains in LPS,^40,41^ did not severely impair the activity of GM. In addition, mutation of mMD-2_F126, which blocks LPS-induced TLR4 dimerization,^40,41^ abolished NF-*κ*B activation by both GM and LPS, suggesting that dimerization of the TLR4/MD-2 complex is required for GM-induced signal transduction. While our study elucidated several key molecular interactions in GM recognition, a detailed understanding of how the disaccharide GM engages with the TLR4/MD-2 complex requires further structural investigations.

### Structure-activity-relationship (SAR) analysis of GM-induced TLR4 signaling

Given the mild stimulation by GM and its weak affinity for TLR4, we questioned the specificity of TLR4 recognition for disaccharide ligands. With HEK-Blue^TM^ mTLR4 reporter cells, we tested a panel of synthetic disaccharides bearing various structural modifications, including deacetylation of GlcNAc (A1), alterations of the lactoyl moiety on MurNAc (A2-A4), 1,6-anhydro-terminus (G-ahM), and the regioisomer MG, as well as chitobiose and diacetylated chitobiose of the natural fungal cell wall (**Fig. 5D**). Remarkably, only GM and MG stimulate TLR4, while other minor structural changes render the disaccharide inactive (**Fig. 5E**), suggesting that TLR4 recognition of gut microbiota-derived disaccharides is highly specific. Furthermore, we showed GM exhibits antagonistic effects against LPS-stimulated TLR4 signaling in the reporter cell, likely by competitively binding to the TLR4 receptor (**Extended Data Fig. 9C**).

### GM mitigates DSS-induced colitis in mice in a TLR4-dependent manner

Upon establishing GM as a bioactive TLR4 ligand *in vitro*, we asked whether this naturally released PGN disaccharide has any biological significance *in vivo*. Given previous work showing that MDP-mediated NOD2 signaling protects against DSS-induced colitis in mice,^42^ we sought to explore the potential efficacy of GM in this context. In our timeline, GM was administered intraperitoneally to mice daily for two days before and throughout a week during which 3% DSS was introduced into their drinking water. For control groups, mice were either injected with GM alone (i.e., no DSS treatment) or PBS buffer (i.e., no GM administration) followed by 3% DSS in their drinking water (**Fig. 6A**). Importantly, we validated that intraperitoneal injection of GM temporally increased its abundance in the mouse gut, providing grounds for daily injections to maintain elevated levels of gut PGN saccharides (**Extended Data Fig. S10A**). The administration of GM (without DSS treatment) did not result in any morbidity in mice. More gratifyingly, GM effectively alleviated DSS-induced colitis in WT mice, as evidenced by reduced body weight loss and colonic shortening (**Fig. 6B-D**). In H&E-stained colonic tissue images, the GM-treated DSS group showed reduced epithelial erosion and mononuclear cell infiltration (**Fig. 6E**, **Extended Data Fig. 10B**). Correspondingly, infiltrating CD45^+^ immune cells, including monocytes and neutrophils, in the colonic lamina propria were significantly reduced in GM-treated DSS mice (**Fig. 6F, Extended Data Fig. 10C).** Consistently, colonic expressions of proinflammatory genes such as *Tnfa*, *Il1b,* and *Ccl2* were significantly suppressed, supporting the effects of GM in attenuating colonic inflammations (**Fig. 6G**). On the other hand, cytoprotective factors of colonic damage such as *keap1* and *hspb1* were upregulated in GM-treated colitis mice (**Extended Data Fig. 10D**). To further assess systemic inflammation in mice, we conducted multiplex immunoassays to quantify 13 cytokines and chemokines in sera of the four groups of WT mice at the timeline endpoint (**Extended Data Fig. 10E**). DSS-induced colitis mice manifested a significant global elevation of inflammatory markers, including IL-6, IFN-*β*, IL-27, IL-10, and IL-23. Remarkably, GM supplementation restored circulating proinflammatory markers to baseline levels, consistent with its ability to mitigate DSS-induced inflammations in mice. On the other hand, GM alone does not significantly alter circulating cytokines and chemokines in mice.

**Figure 6.**
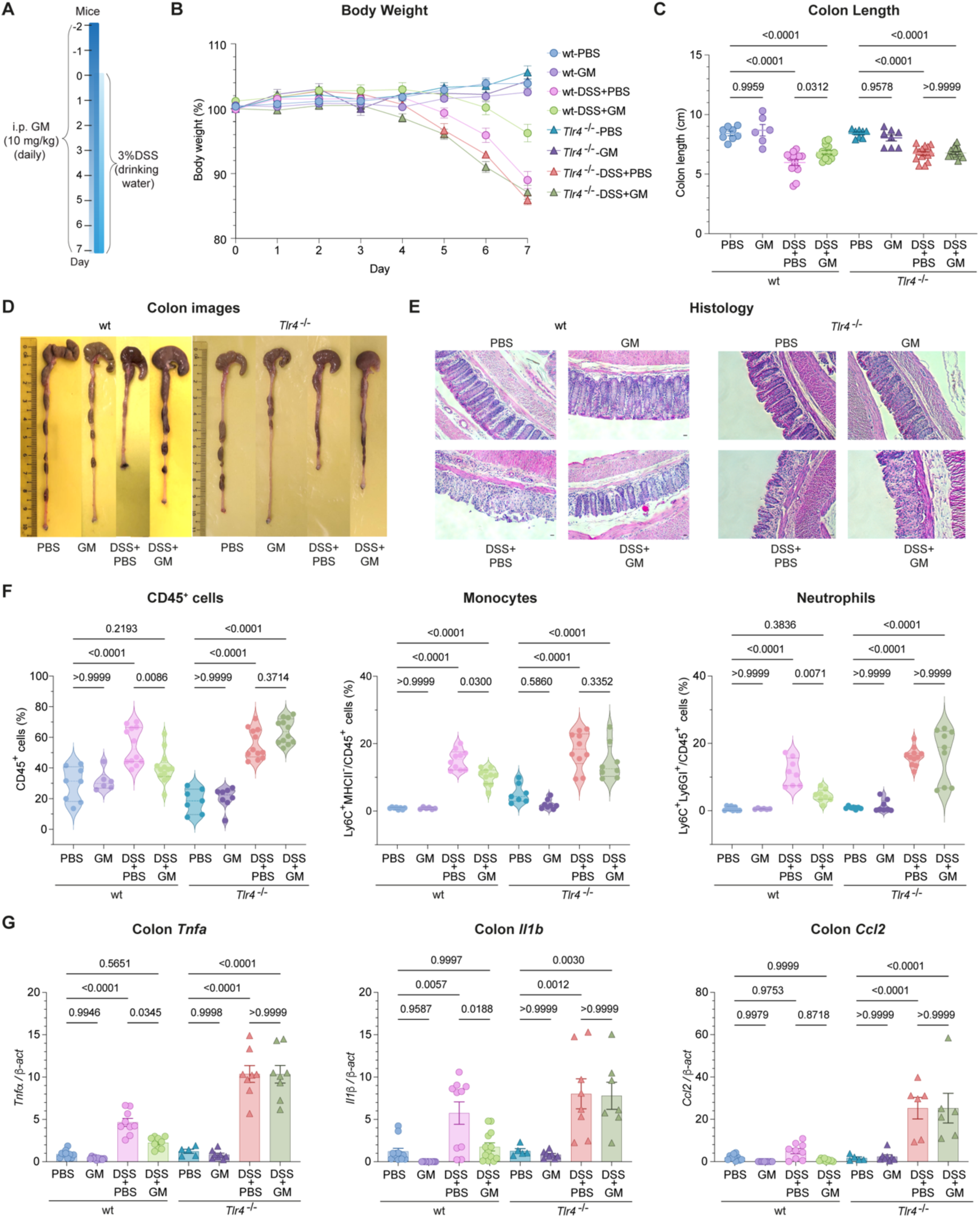
Administration of GM reduces colon inflammation in DSS-induced colitis in WT mice but not *Tlr4*^-/-^ mice. **(A)** Timeline of the study, where GM was intraperitoneally administered at 10 mg/kg to mice (WT and *Tlr4*^-/-^) that were provided with 3% DSS or normal drinking water (*n* = 6-14). I.p. injections of PBS were given to the negative control group. (**B-C**) Measurements of mice body weight (**B**) and colon length (**C**) at the endpoint of the timeline. Data are presented as mean values ± s.e.m. (*n* = 6-14). Statistical significance was determined using two-way ANOVA. (**D-E**)Representative images of colons **(D)** and H&E-stained colon sections (**E**) from each group. (**F**) Flow cytometry analysis of immune cell infiltration in colons of each treatment group. The percentage of the CD45^+^ population was calculated within pre-gated live cells. Percentage of Ly6C^+^ MHCII^-^ and Ly6C^+^ Ly6G^+^ populations were calculated within the CD45^+^ population. Data are presented as mean values ± s.e.m. (*n* = 6-14). Statistical significance was determined using two-way ANOVA. (**G**) RT-qPCR analysis of colonic cytokines demonstrates the significantly reduced inflammation in the GM-supplemented DSS-induced colitis in WT mice but not *Tlr4^-/-^* mice. Data are presented as mean values ± s.e.m. (*n* = 6-10). Statistical significance was determined using two-way ANOVA.

To determine whether GM’s *in vivo* protective effects are TLR4-dependent, we performed the aforementioned experiments in *Tlr4^-/-^* mice (**Fig. 6A**). In this case, GM administration failed to prevent weight loss and colonic shortening in DSS-administered *Tlr4^-/-^* mice (**Fig. 6B-D**). In addition, similar levels of colonic crypt damage were observed in the colitis *Tlr4*^-/-^ mice, regardless of GM supplementation (**Fig. 6E**). The results of immune cell infiltration and colonic proinflammatory gene expression were consistent with the loss of GM’s protective effects against DSS-induced colitis in *Tlr4^-/-^* mice (**Fig. 6G)**. No upregulation of cytoprotective responses in GM-treated colitis *Tlr4^-/-^* mice was observed (**Extended Data Fig. 10D**). Collectively, our data suggest that the gut microbiota-derived disaccharide GM maintains host gut homeostasis via TLR4-dependent mechanisms.

## Discussion

Using the LC-MS/MS-based PGN analysis platform, we discovered that 90% of the gut microbiota-derived PGNs in the host gut consist solely of saccharide moieties, while the remaining muropeptides are primarily of the mDAP type. Given that the gut bacterial peptidoglycome encompasses both Lys- and mDAP-type peptidoglycan from Gram-positive and Gram-negative bacteria,^14^ it is intriguing that the soluble muropeptides in the host gut milieu appear mostly derived from Gram-negative gut bacteria. We speculated that gut commensals such as *Bacteroides spp*. that possess mDAP-containing peptidoglycan could be the major contributors to gut PGNs.^43^ Moreover, we did not identify the model ligand MDP (M-AQ) in hosts; instead, we found that its non-amidated analogues, such as M-AE and GM-AE, are prevalent in the mice ceca. These natural muramyl dipeptides are likely the cleavage products of commensal bacteria-secreted peptidoglycan endopeptidases, such as SagA and

LPH. Previous studies have shown that elevating the levels of these enzymes or their corresponding cleaved muropeptide products benefits gut homeostasis and enhances cancer immunotherapy in mice.^7,44–46^ On the other hand, we only detected a minute amount of PGN monosaccharide (e.g. M and ahM) in healthy host sera, which could indicate selective systemic dissemination of gut PGNs^25^ or PGN degradation by serum amidase PGRP2.^31^ Alternatively, circulating PGNs may be retained by serum proteins or carried by vesicular transporters,^25,47,48^ potentially evading detection by LC-MS/MS. Nevertheless, we emphasize that, unlike other existing PGN detection assays that rely on protein recognition of specific motifs, our LC-HRMS/MS analysis is unbiased and directly identifies PGN structures in hosts.

Intrigued by the abundance of PGN saccharides that do not resemble classic NOD1/2 ligands in the host gut, we were prompted to investigate the potential bioactivity of these non-canonical PGN moieties. Wolf *et al.* previously reported that cytosolic GlcNAc released from the phagosome degradation of bacterial peptidoglycan is detected by hexokinase that leads to inflammasome activation in LPS-primed immune cells, unveiling a NOD1/2-independent PGN sensing mechanism in the host.^21^ Conversely, MurNAc was also shown to suppress inflammation in LPS-induced macrophages *in vitro*.^49^ Herein, we demonstrated that the gut microbiota-derived disaccharide GM exerts immuno-stimulatory effects in the absence of LPS priming, under which condition the monosaccharides G and M are inactive. Importantly, we established that GM acts as a TLR4 agonist, which directly binds to TLR4 and activates downstream NF-κB and IRF pathways. Of note, bacterial LPS is the canonical TLR4 ligand, whose Lipid A motif constitutes a *β*-1,6-linked glucosamine disaccharide with phosphate groups and multiple acyl chains.^50^ Apart from LPS, endogenous ligands such as heat shock proteins, hyaluronan, and monosodium urate crystals, as well as synthetic small molecules like neoseptins are reported to stimulate TLR4 signaling, demonstrating the structural diversity of TLR4 ligands.^51–53^ Our discovery of the gut microbiota-derived disaccharide GM as a TLR4 agonist expands the scope of natural TLR4 ligands in hosts. Structurally, GM may resemble the disaccharide core of Lipid A for TLR4 recognition; however, the lack of hydrophobic acyl chains in GM renders it a mild TLR4 agonist. The fact that TLR4 selectively recognizes disaccharide PGN motifs (i.e., GM and its regioisomer MG) but not closely related fungal cell wall fragments underscore the potential biological significance of natural PGNs in gut microbiota-host interactions.

Notably, TLR4 activation by metabolites from gut commensal microflora under steady-state conditions is essential for maintaining gut homeostasis and protecting against colonic injury.^54^ For instance, LPS supplementation effectively rescues the severity of DSS-induced colitis in commensal-depleted mice via TLR4 stimulation.^54,55^ Our discovery of the naturally abundant gut microbiota-derived disaccharide GM as a novel TLR4 ligand suggests its physiological relevance in hosts. Indeed, the administration of GM protects against DSS-induced colitis in mice via TLR4-dependent mechanism(s). We offer several hypotheses accounting for its *in vivo* protection. First, GM, as a mild TLR4 agonist, may act to prime TLR4 to downregulate its inflammatory responses to subsequent triggers in hosts. Supportively, we observed that GM antagonizes LPS-induced activation of TLR4-mediated NF-κB signaling in reporter cells *in vitro*. Second, colonic epithelium TLR4 activation confers cytoprotection and repair against DSS-induced tissue injury and damage.^54^ To this end, we confirmed that GM supplementation indeed upregulated cytoprotective genes such as *keap1* and *hspb1* in colonic tissues of the DSS-treated wildtype but not *Tlr4*^-/-^ mice. Lastly, GM-induced TLR4 activation may in turn regulate other signaling pathways to provide protective effects *in vivo*. For example, MDP activation of NOD2 suppresses DSS-induced colitis in mice via the induction of negative regulator IRF4.^42^ Intriguingly, our transcriptomics analysis in this work uncovered the intricate effects of TLR4 on MDP-mediated NOD2 signaling, as indicated by the reduced expressions of specific genes in MDP-treated *Tlr4*^-/-^ BMDMs compared to treated WT BMDMs. Extrapolating from these *in vitro* observations, we hypothesize that GM supplementation stimulates TLR4 signaling, which may promote NOD2 activation by endogenous gut microbiota-derived muropeptides like M-AE and GM-AE to protect against DSS-induced colitis in mice. The in-depth mechanisms will be explored in future studies. Importantly, from a therapeutic perspective, the gut microbiota-derived disaccharide GM—a mild TLR4 agonist that does not induce acute inflammation or morbidity in mice—emerges as a promising candidate for postbiotics or adjuvants. It holds attractive potential for enhancing TLR4-mediated immune responses against gut-associated inflammatory diseases.

## Acknowledgment

We acknowledge Prof Yue Wang and Ms Xiaoli Xu (A*STAR Infectious Disease Labs) for providing 2E7 monoclonal antibody, SPF mice fecal and cecal samples for initial analysis, as well as for helpful discussions throughout the project. We acknowledge Dr. Hwei Ee Tan (A*STAR and NTU) for providing GF mice fecal samples. We thank Effie Lee and Wilaiporn Saikruang for assistance with the study. The work was supported by the National Research Foundation (NRF) Singapore, NRF-NRFF12-2020-0006, Nanyang Technological University Start-up grant (SUG), and College of Science Collaborative Research Award 2021 to Y.Q.

## Author contributions

Conceptualization, C.L. and Y.Q.; Methodology, C.L., J.T.L., S.K.N., C.R., S.H.W., K.P.L., and Y.Q.; Investigation, C.L., C.A., A.W.R.N., Y.L., Z.H., J.T.L., S.K.N., S.F., E.W.L.N., N.S.K., and E.L.; Formal analysis, C.L., J.K.M.C., and Y.Q.; Writing – Original Draft, C.L. and Y.Q.; Writing – Review & Editing, C.L., C.A., A.W.R.N., J.L., J.M.C.K., C.R., S.H.W., K.P.L., and Y.Q.; Resources, C.R., S.H.W., K.P.L. and Y.Q.; Funding Acquisition and Supervision, Y.Q.

